# The influence of generativity on purpose in life is mediated by social support and moderated by prefrontal functional connectivity in at-risk older adults

**DOI:** 10.1101/2023.02.26.530089

**Authors:** Caitlin S. Walker, Linda Li, Giulia Baracchini, Jennifer Tremblay-Mercier, R. Nathan Spreng, The PREVENT-AD Research Group, Maiya R. Geddes

## Abstract

**Objectives:** Generativity, the desire and action to improve the well-being of younger generations, is positively associated with purpose in life among older adults. However, the neural basis of generativity and the neurobehavioral factors supporting the relationship between generativity and purpose in life remain unknown. This study aims to identify the functional neuroanatomy of generativity and mechanisms linking generativity with purpose in life in at-risk older adults.

**Methods:** Fifty-eight cognitively healthy older adults (mean age = 70.78, 45 females) with a family history of Alzheimer’s disease were recruited from the PREVENT-AD aging cohort. Participants underwent brain imaging and completed questionnaires assessing generativity, social support, and purpose in life. Mediation models examined whether social support mediated the association between generativity and purpose in life. Seed-to-voxel analyses investigated the association between resting-state functional connectivity (rsFC) to the ventromedial prefrontal cortex (vmPFC) and ventral striatum (VS) and whether this rsFC moderated the relationship between generativity and purpose in life.

**Results:** Affectionate social support mediated the association between generative desire and purpose in life. Generative desire was associated with rsFC between VS and precuneus and vmPFC and right dorsolateral prefrontal cortex (rdlPFC). The vmPFC-rdlPFC connectivity moderated the association between generative desire and purpose in life.

**Discussion:** These findings provide insight into how the brain supports social behavior and, separately, purpose in life in at-risk aging. Affectionate social support may be a putative target process to enhance purpose and life in older adults. This knowledge contributes to future developments of personalized interventions that promote healthy aging.

## 1. Introduction

Alzheimer’s disease (AD) is a progressive neurodegenerative disease characterized by a decline in cognitive abilities, such as memory, learning, and reasoning (DeTure & Dickson, 2019). Identifying disease prevention strategies is crucial, as delaying the onset of AD by 5 years could reduce its prevalence by 50% (Brookmeyer et al, 1998). One significant risk factor associated with a 1.26x increased risk of dementia is social isolation (Shen et al, 2022). Older adults who have high levels of perceived social support show higher cognitive abilities and a reduced risk of dementia, even in cases of preclinical AD or mild cognitive impairment (Marioni et al., 2015). This suggests that social support plays a vital role in preventing AD-related cognitive decline.

There is growing evidence that older adults undergo a ‘prosocial goal shift’ where behaviours aimed at benefitting the well-being of others become increasingly rewarding with age (Isaacowitz, 2021). A key aspect of social motivation is generativity, which involves both the desire to contribute to the well-being of younger generations (referred to as generative *desire*) and actions directed towards increasing their well-being (referred to as generative *achievement;* Gruenewald et al., 2016). Originally defined by Erikson (1950) in his psychosocial development theory, generativity was described as a stage at midlife that arises from a desire to be needed. Erikson proposed that generativity in midlife evolves into “grand-generativity” during older adulthood characterized by a reflective evaluation of one’s life and the legacy they leave behind. Prior studies have shown that generativity remains stable or even increases in older adulthood (Isaacowitz, 2021). Moreover, it is associated with life satisfaction, well-being, participation in productive activities, self-efficacy, and physical health (Gruenewald et al, 2012; Pinazo-Hernandis et al., 2023).

The relationship between generativity and well-being in older adulthood aligns with the predictions of Socioemotional Selectivity Theory (SST; Carstensen, 2006). SST posits that an individual’s perception of their time horizons influences their prioritization of social goals (Carstensen, 2006). In young adulthood when time is perceived as open-ended, goals involving knowledge acquisition are prioritized. In contrast, in older adulthood when time is perceived as limited, goals that enhance socioemotional well-being in the present take precedence. Generativity, by promoting the well-being of future generations, can serve as a means for older adults to maximize their socioemotional well-being. Notably, SST seems to apply to older adults in the preclinical and early stages of AD where mental states that promote prosocial behavior, including positive affect, social attention, and empathic concern, are preserved or enhanced compared to age-matched controls (Chow et al., 2023; Sturm et al., 2004). These findings raise the intriguing possibility that social factors may be particularly important intervention targets for AD prevention.

Generativity can be enhanced in older adulthood through volunteer work that establishes intergenerational connections. For instance, the Baltimore Experience Corps Trial (BECT) is an intergenerational social program where older adults volunteer to help elementary school students with reading. Compared to a control group, participants in this program showed significant increases in generative desire and achievement following the intervention, and this effect was sustained after 24 months (Gruenewald et al., 2016). Additionally, older adults who participated in the program demonstrated improved physical, social, and cognitive functioning compared to a control group (Carlson et al., 2009; Tan et al., 2006). A subgroup of older adults in this study who were at the greatest risk of cognitive impairment (i.e., Mini-Mental State Examination score < 24 or diminished executive function at baseline) showed the greatest improvement in memory, executive function and prefrontal activation following the intervention (Carlson et al., 2009; Carlson et al., 2008). This suggests that intergenerational programs promoting generativity might be particularly effective for older adults at a greater risk of cognitive decline.

Converging behavioral studies have demonstrated the positive association between generativity and purpose in life in older adults (de St. Aubin, 2013). Purpose in life is conceptualized as having goals, directedness, and feelings that give one’s life meaning (Boyle et al., 2010). According to McAdams and de St. Aubin’s (1992) generativity model, the integration of generativity into older adults’ lives allows them to construct a meaningful identity and life story, leading to purpose in life (Kruse & Schmitt, 2012). This life purpose is associated with many facets of successful aging, including increased physical health, greater engagement in health behaviors, and a reduced risk of AD and dementia (Boyle et al., 2010; Pinquart, 2002).

Older adults with AD or dementia reveal that their family roles and relationships hold the most value to them (Cohen-Mansfield et al., 2006; Cotrell & Hooker, 2005). These social connections are believed to motivate older adults to engage in various activities and interests, which can enhance purpose in life (Pinquart, 2002). Moreover, older adults who exhibit high levels of generativity are more likely to report greater social support than those with lower generativity (Hart et al., 2001), and interventions designed to enhance generativity have also demonstrated increases in perceived social support (Moieni et al., 2021). Despite these independent observations, prior research has yet to examine if higher social support through enhanced generativity improves older adults’ purpose in life.

Understanding the neural mechanisms supporting generativity is a crucial step in comprehending and potentially modifying this behaviour. To date, no studies have examined the functional neuroanatomy of generativity, making it an important area of research for developing brain-based interventions to promote health and resilient aging. Resting-state functional connectivity (rsFC) is a powerful method for studying the functional organization of the brain and the networks supporting internal mental processes and behavior (Stevens & Spreng, 2014). Based on prior research, it is possible that generativity relies on brain systems implicated at the intersection of value-based decision making and self-transcendence. A coordinate-based meta-analysis identified the ventromedial prefrontal cortex (vmPFC) and ventral striatum (VS) as nodes of the reward network implicated in value-based decision making by representing the subjective value of decision options (Bartra, 2013; Zhang & Glascher, 2020). The vmPFC is particularly important in considering emotional content during subjective valuation and demonstrates higher activity when older adults attend to positive relative to negative stimuli (Mather, 2016; Vaidya et al., 2018). Rademacher et al. (2014) found that there was enhanced activity in the VS for social compared to monetary rewards for older adults, while activity in the VS was higher for monetary compared to social rewards for younger adults. These results highlight the potential involvement of the vmPFC and VS in social motivation, including generativity. Moreover, self-transcendence, the process of shifting one’s focus from self-interests to the well-being of others and humanity, is closely related to generativity. Previous research has shown that self-transcendence and prosocial behaviour are associated with activity in the vmPFC and VS (Kang et al., 2018; Marsh, 2016). As intergenerational social engagement tends to be emotionally rewarding and promotes purpose for older adults (Isaacowitz, 2021; Lyndon & Moss, 2022), differential connectivity of the vmPFC and VS may enhance the relationship between generativity and purpose in life.

This study has two primary aims. The first aim is to identify a behavioral mechanism that explains how generativity enhances purpose in life in at-risk older adults. First, it is hypothesized that generativity will be positively associated with purpose in life, thus replicating prior findings in a sample of older adults at risk of AD. Given that social support is associated with both generativity and purpose in life in older adults, it is hypothesized that social support will mediate the relationship between generativity and purpose in life. The second aim of this study is to identify the functional neuroanatomy supporting generativity and purpose in life in older adults at risk of AD. It is hypothesized that generativity will be positively associated with rsFC to the vmPFC and VS, key regions at the convergence of reward valuation and self-transcendence. It is additionally hypothesized that rsFC to the vmPFC and VS will moderate the association between generativity and purpose in life, providing insight into the neural mechanisms influencing the relationship between generativity and purpose in life.

## 2. Methods

### 2.1 Participants

Participants consisted of 58 cognitively normal older adults (*M*_age_ = 70.78, 45 female) from the PResymptomatic EValuation of Experimental or Novel Treatments for Alzheimer’s Disease (PREVENT-AD) cohort at McGill University (Tremblay-Mercier et al., 2021). A Monte Carlo power analysis (Schoemann et al., 2017) determined that a sample size of *N* = 58 was required for detecting an indirect effect with power of 80%, correlations of .40 for the *a* and *c’* paths, and a correlation of .50 for the *b* path. Fifty participants from the original sample had MRI data available and were included in the rsFC analyses. G*Power version 3.1 (Faul et al., 2007) was used to conduct power analyses for the moderation analysis. The analysis showed that a sample size of *N* = 50 was required for detecting an interaction with a small-to-medium effect size of *f^2^* = 0.13 and power of 80%.

The PREVENT-AD cohort consists of adults 55 years and older who are cognitively normal at the time of enrollment and have a first-degree family history of AD. Participants were considered *APOE ε4* carriers if they had at least one *ε4* allele. Exclusion criteria included history of a major psychiatric illness (e.g., schizophrenia), current treatment for cancer (excluding non-melanoma skin cancer), diagnosis of a neurological disorder or brain injury, current alcohol or substance abuse, a cardiac event that occurred in the past six months (e.g., myocardial infarction, coronary artery bypass grafting, angioplasty) or any change in an antipsychotic, anti-depressant, anti-anxiety, or attention deficit disorder / attention deficit hyperactivity disorder medication in the past six months. Participants’ demographic characteristics are shown in Table 1.

**Table 1.**
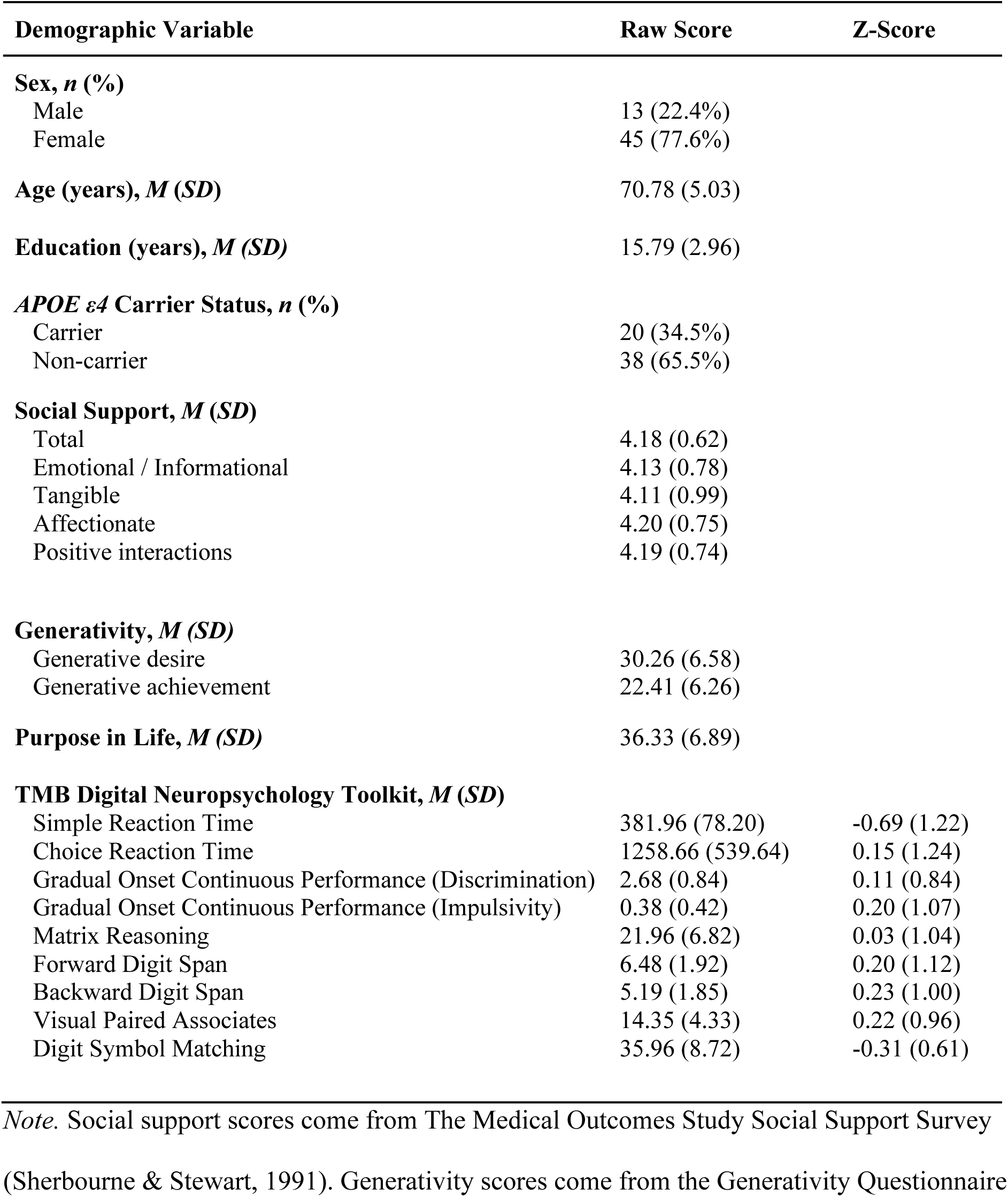
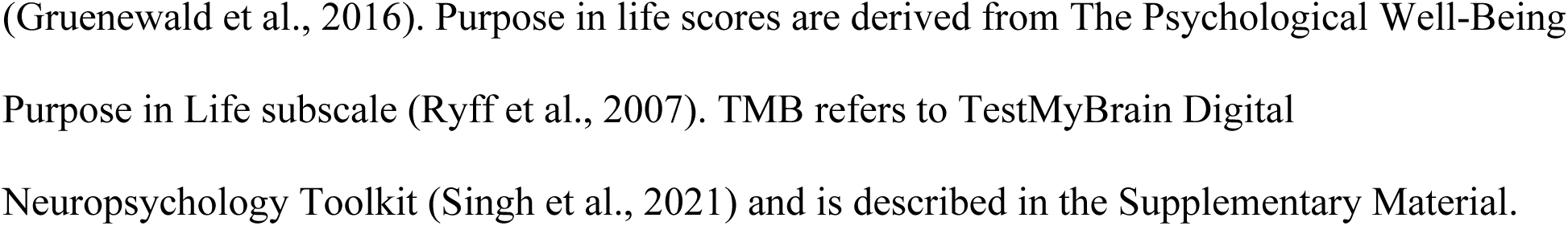
Demographics, neurobehavioral inventory scores, and computerized cognitive assessment scores for the full sample (N = 58)

| Demographic Variable | Raw Score | Z-Score |
| --- | --- | --- |
| **Sex, *n* (%)** |  |  |
| Male | 13 (22.4%) |  |
| Female | 45 (77.6%) |  |
| **Age (years), *M* (*SD*)** | 70.78 (5.03) |  |
| **Education (years), *M* (*SD*)** | 15.79 (2.96) |  |
| ***APOE* ε4 Carrier Status, *n* (%)** |  |  |
| Carrier | 20 (34.5%) |  |
| Non-carrier | 38 (65.5%) |  |
| **Social Support, *M* (*SD*)** |  |  |
| Total | 4.18 (0.62) |  |
| Emotional / Informational | 4.13 (0.78) |  |
| Tangible | 4.11 (0.99) |  |
| Affectionate | 4.20 (0.75) |  |
| Positive interactions | 4.19 (0.74) |  |
| **Generativity, *M* (*SD*)** |  |  |
| Generative desire | 30.26 (6.58) |  |
| Generative achievement | 22.41 (6.26) |  |
| **Purpose in Life, *M* (*SD*)** | 36.33 (6.89) |  |
| **TMB Digital Neuropsychology Toolkit, *M* (*SD*)** |  |  |
| Simple Reaction Time | 381.96 (78.20) | -0.69 (1.22) |
| Choice Reaction Time | 1258.66 (539.64) | 0.15 (1.24) |
| Gradual Onset Continuous Performance (Discrimination) | 2.68 (0.84) | 0.11 (0.84) |
| Gradual Onset Continuous Performance (Impulsivity) | 0.38 (0.42) | 0.20 (1.07) |
| Matrix Reasoning | 21.96 (6.82) | 0.03 (1.04) |
| Forward Digit Span | 6.48 (1.92) | 0.20 (1.12) |
| Backward Digit Span | 5.19 (1.85) | 0.23 (1.00) |
| Visual Paired Associates | 14.35 (4.33) | 0.22 (0.96) |
| Digit Symbol Matching | 35.96 (8.72) | -0.31 (0.61) |

*Note.* Social support scores come from The Medical Outcomes Study Social Support Survey
(Sherbourne & Stewart, 1991). Generativity scores come from the Generativity Questionnaire
(Gruenewald et al., 2016). Purpose in life scores are derived from The Psychological Well-Being Purpose in Life subscale (Ryff et al., 2007). TMB refers to TestMyBrain Digital Neuropsychology Toolkit (Singh et al., 2021) and is described in the Supplementary Material.

### 2.2 Procedure

All study procedures were approved by the McGill University Research Ethics Board. Participants from the PREVENT-AD cohort were recruited for the study by contacting them via phone and email and by sending flyers in the mail. All eligible participants then attended a videoconferenced meeting with a member of the research team where they were provided with detailed information about study procedures and were given the opportunity to ask any questions. All participants provided written informed consent in accordance with the Declaration of Helsinki. Thereafter, all participants completed an online battery of questionnaires, described below.

### 2.3 Behavioral Inventories

#### The BECT Generativity Questionnaire (Gruenewald et al., 2016)

This measure was used to assess generative desire and achievement. Participants rated the extent to which they agree with seven generative desire statements (e.g., “I want to make a difference in the lives of others.”) and six generative achievement statements (e.g., “I feel like I make a difference in my community.”) on a 6-point scale ranging from *strongly disagree* to *strongly agree*. Scores for the generative desire subscale ranged between 7 and 42, with higher scores indicating greater generative desire. Scores for the generative achievement subscale ranged between 6 and 36, with higher scores indicating greater generative achievement.

#### The Medical Outcomes Study Social Support Survey (Sherbourne & Stewart, 1991)

This measure consists of 19 items that assess the availability of four types of social support. The emotional and informational support subscale consists of eight items that measure how often individuals have someone to listen to them, confide in, and provide them with advice and information. The tangible support subscale consists of four items that measure the frequency of support in daily tasks such as chores and preparing meals. The affectionate support subscale consists of three items that measure how often individuals have someone to show them love and affection. Finally, the positive social interaction subscale consists of three items that measure how often individuals have someone to spend time and do enjoyable things with. Each item is scored on a Likert scale of 1 to 5 where 1 = *none of the time* and 5 = *all the time*. Scores for each subscale are obtained by calculating the mean score across its items. All scores range between 1 and 5, with higher scores indicating a higher availability of social support.

#### The Psychological Well-Being Purpose in Life subscale (Ryff et al., 2007)

This inventory consists of seven items that measure the extent to which individuals have goals, a sense of directedness, and beliefs that give life meaning (e.g., “Some people wander aimlessly through life, but I am not one of them.”). Each item is scored using a Likert scale from 1 to 7 where 1 = *strongly disagree* and 7 = *strongly agree*. Negatively worded items are reverse coded, then scores on all items are summed to create a total score. Total scores range between 7 and 49, with higher scores indicating greater purpose in life.

### 2.4 MRI Data Acquisition

Participants were scanned in a 3T Siemens Magnetom Prisma MRI scanner using a standard Siemens 32-channel head coil (Siemens Medical Solutions, Erlangen, Germany). T1-weighted structural images were acquired using an MPRAGE (Magnetization Prepared Rapid Gradient Echo Imaging) sequence. The parameters included a repetition time (TR) of 2300 milliseconds (ms), echo time (TE) of 2.96 ms, inversion time (TI) of 900 ms, flip angle of 9°, Field of View (FOV) of 256 millimeters (mm), phase encode A-P, a GRAPPA acceleration factor of 2, and a Bandwidth (BW) of 625 Hz/px. The rsfMRI data were acquired by an echo-planar imaging (EPI) sequence. The scanning duration for each run was 5.04 minutes and two sessions were performed continuously. The parameters were as follows: TR = 1000 ms, TE1 = 12 ms, TE2 = 30.11 ms, TE3 = 48.22 ms, flip angle = 50°, FOV = 240 mm, phase encode A-P, BW = 2500 Hz/px. Forty-eight slices were collected in each run. Information on the preprocessing and denoising of the functional and structural data can be found in the Supplementary Materials.

### 2.5 Analyses

#### 2.5.1 Behavioral Analyses

First, two linear regression models were conducted to determine whether generative desire and achievement were associated with purpose in life. Then, Model 4 of the PROCESS macro for SPSS, version 4.2 (Hayes, 2022) was used for the mediation analyses. Generative desire and generative achievement were entered as predictor variables in separate models, with purpose in life included as the outcome variable in each. Sex, age, and *APOE ε4* carrier status were included as covariates in all mediation models. Affectionate social support and positive social interactions scores were significantly correlated with generative desire, generative achievement, and purpose in life (*p*s < .05). Therefore, four separate models were computed with affectionate social support and positive social interactions included as mediators and generative desire and generative achievement included as predictors of purpose in life. The indirect effects were tested using a percentile bootstrapping method with 1000 samples. Because four separate mediation models were computed, we applied a Bonferroni correction and used 99% confidence intervals for detecting significant indirect effects.

#### 2.5.2 Functional Brain Imaging Analyses

Seed-to-voxel rsFC analyses were performed using the CONN toolbox (Whitfield-Gabrieli & Nieto-Castanon, 2012). Two regions of interest (ROIs) in the vmPFC and VS were functionally defined based on masks from Batra et al.’s (2013) meta-analytic conjunction analysis on the neural correlates of subjective value. The average time series in the vmPFC and VS ROIs were extracted. First-level correlation maps were obtained by extracting product-moment correlation coefficients between the average time series in each ROI and the time series of all other voxels. Correlation coefficients were then Fisher transformed to z-scores to increase normality prior to the second-level general linear models. Seed-to-voxel maps for the vmPFC and VS are included in Figures 1 and 2 of the Supplementary Materials. Two general linear models were computed for each ROI, with generative desire and generative achievement included as the covariates of interest. Age, sex, mean head framewise displacement, and *APOE ε4* carrier status were included as nuisance variables in the general linear models. All analyses were performed with voxel level *p* < .001 and a cluster-size *p* < .05 family wise error (FWE) corrected cluster-extent threshold to correct for multiple comparisons.

**Figure 1.**
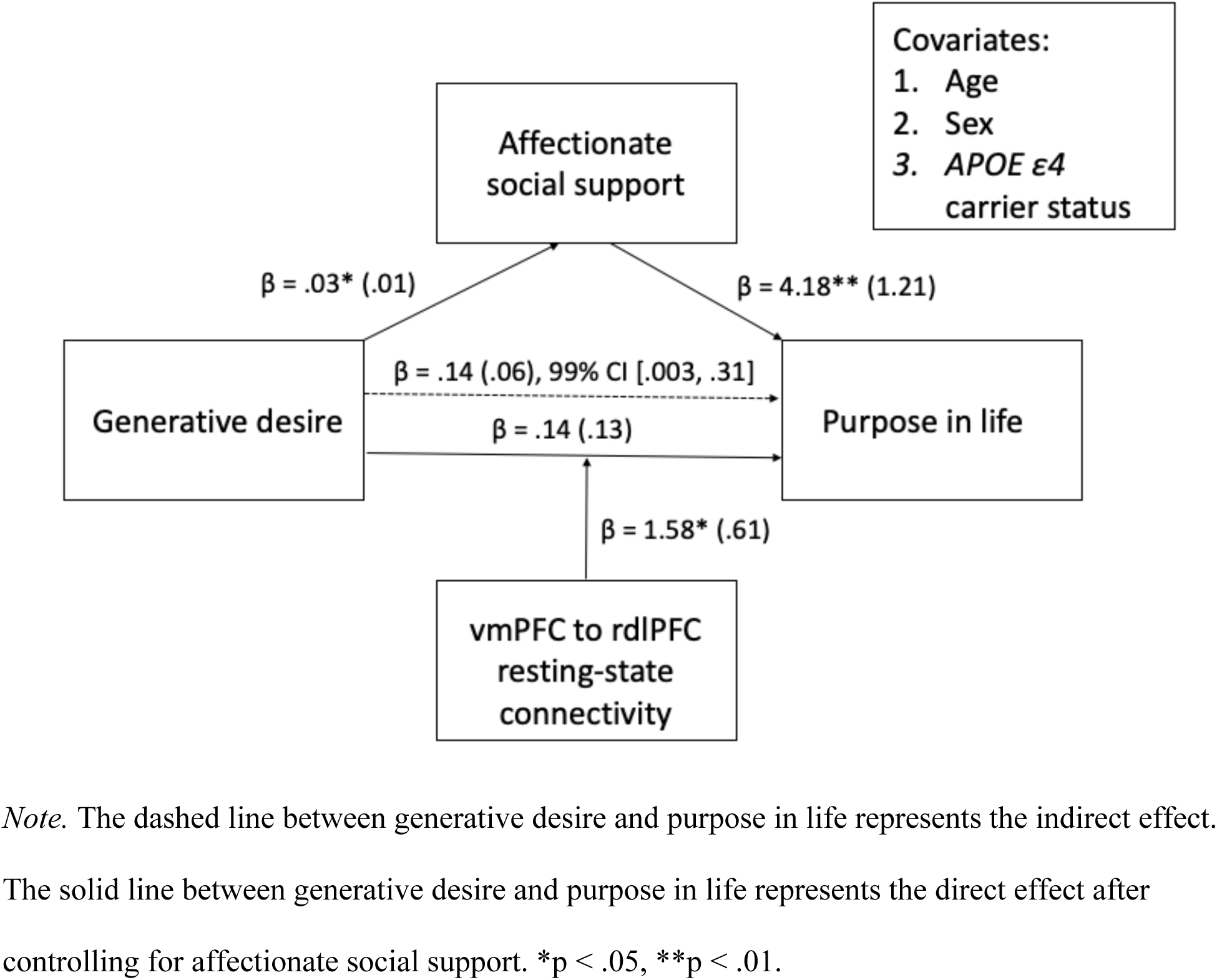
Diagram of the Mediation and Moderation Model Results Between Generative Desire and Purpose in Life.

**Figure 2.**
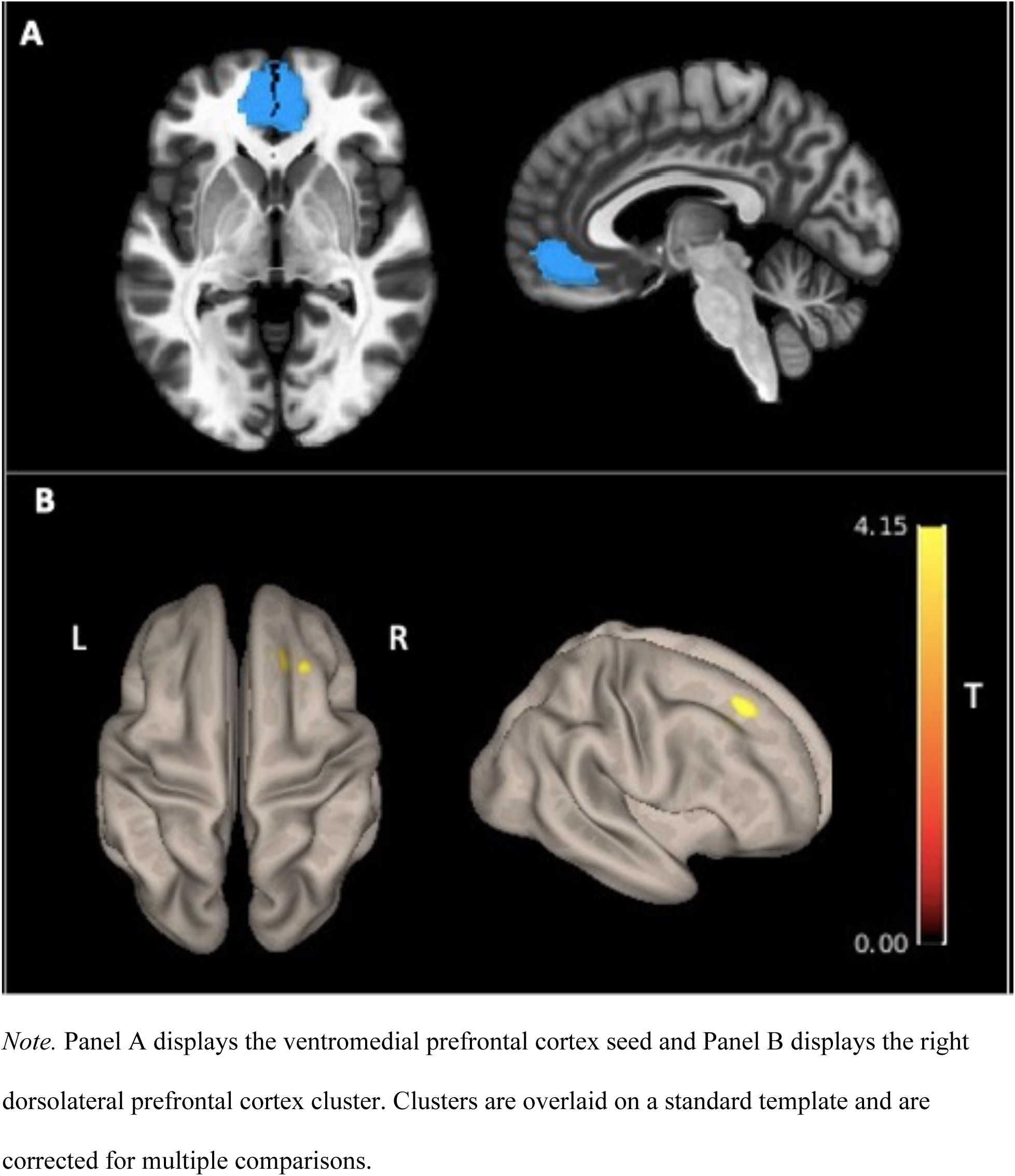
Resting-State Functional Connectivity Between the Ventromedial Prefrontal Cortex and Right Dorsolateral Prefrontal Cortex is Associated with Generative Desire.

Next, a moderation analysis using Model 1 of the SPSS PROCESS macro was conducted to examine the interactive effect between rsFC and generative desire on purpose in life. Fisher-transformed z-scores were extracted from the rsFC analyses. Generative desire and rsFC values were mean centered to reduce multicollinearity. A generative desire x rsFC interaction term was computed and entered as a predictor of purpose of line in the regression models.

## 3. Results

### 3.1 Behavioral

Separate linear regression models revealed that generative desire (β = 0.29, SE = 0.14, 95% CI [0.01, 0.56], *p* = .040, R^2^ = 0.07) and generative achievement (β = 0.42, SE = 0.14, 95% CI [0.15, 0.69], *p* = .003, R^2^ = 0.15), were significant predictors of purpose in life. The model with affectionate social support included as the mediator between generative desire and purpose in life was statistically significant (Figure 1). In particular, the direct effect of generative desire on purpose in life was no longer statistically significant after controlling for the effect of affectionate social support (β = 0.14, SE = 0.13, 99% CI [-0.21, 0.50], *p* = .276). Furthermore, the indirect effect was statistically significant (β = 0.14, SE = 0.06, bootstrapped 99% CI [0.003, 0.31]). This suggests that affectionate social support fully mediated the effect of generative desire on purpose in life. The mediation models with generative achievement as the predictor or positive social interactions as the mediator variable did not reveal statistically significant indirect effects.

### 3.2 Functional Brain Imaging

As shown in Figure 2, generative desire was associated with enhanced rsFC between the vmPFC seed and a cluster in the right dorsolateral prefrontal cortex (rdlPFC; *t*(45) = 4.43, voxel *p* < 0.001 uncorrected, cluster *p*-FWE < 0.05, peak voxel MNI coordinates = [24, 32, 44], k = 54). Generative desire was also positively associated with rsFC between the VS seed and a cluster in the right precuneus, (Figure 3; *t*(45) = 5.55, voxel *p* < 0.001 uncorrected, cluster *p*-FWE < 0.05, peak voxel MNI coordinates = [06, −58, 54], k = 154). A bivariate product-moment correlation analysis showed that generative desire was not significantly correlated with mean head framewise displacement, *r* = .151, *p* = .333. Seed-to-voxel analyses with generative achievement included as the covariate of interest were not statistically significant.

**Figure 3.**
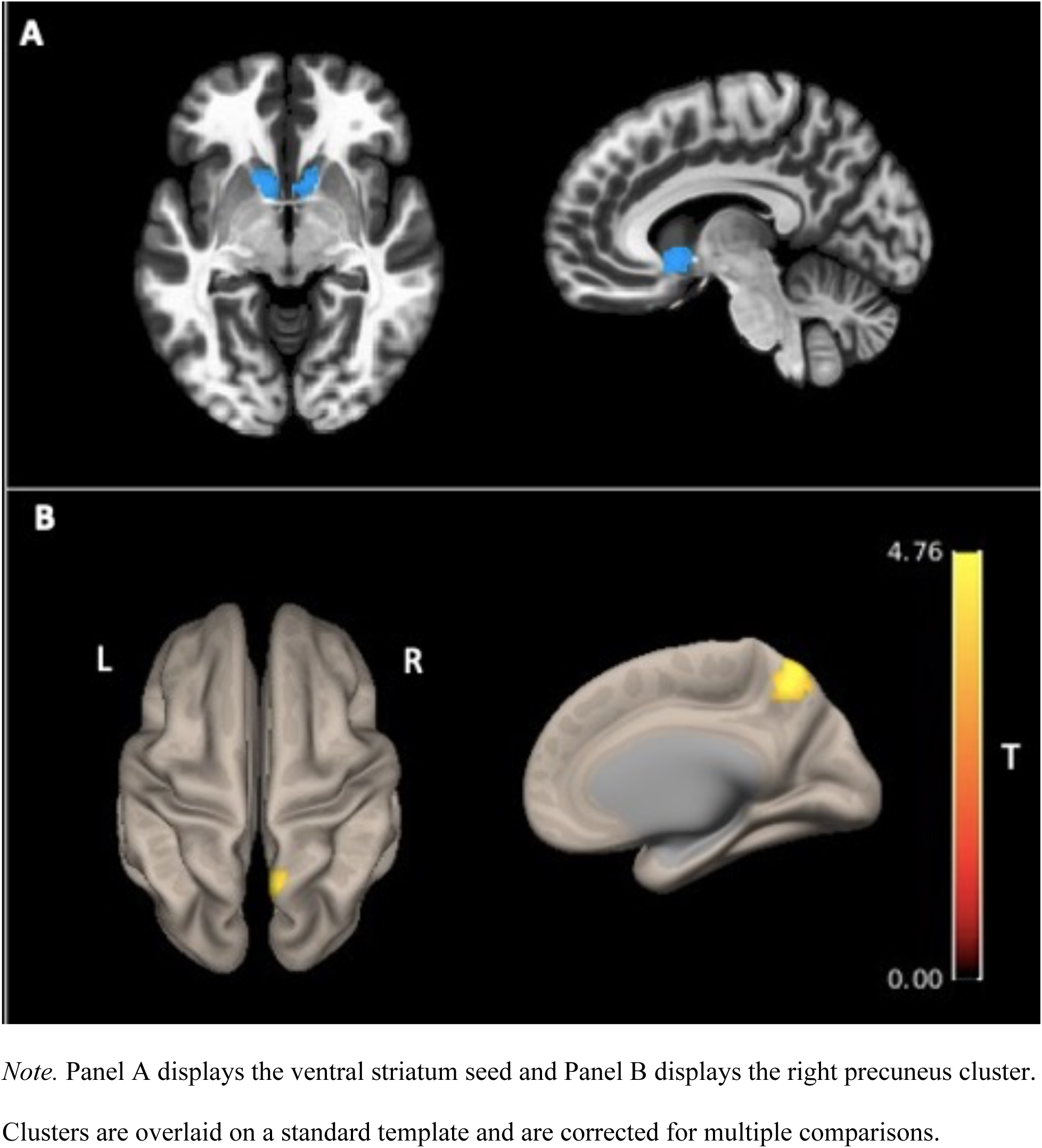
Resting-State Functional Connectivity Between the Ventral Striatum and Right Precuneus is Associated with Generative Desire.

The interaction between generative desire and rsFC between the vmPFC and rdlPFC on purpose in life was also statistically significant (β = 1.58, SE = 0.61, 95% CI [0.36, 2.81], *p =* .013). Simple slopes for the association between generative desire and purpose in life at low (−1 standard deviation (SD) below the mean), moderate (mean), and high (+1 SD above the mean) levels of rsFC are shown in Figure 4. Simple slope tests revealed a statistically significant association between generative desire and purpose in life at high rsFC values (β = 0.70, SE = 0.26, 95% CI [0.18, 1.22], *p =* .010) and a marginally significant association for moderate rsFC values (β = 0.37, SE = 0.19, 95% CI [-0.004, 0.75], *p =* .052). The simple slope was not statistically significant at low rsFC (*p* = .824). The interaction between generative desire and VS-precuneus rsFC approached but failed to reach statistical significance, *p* = .084.

**Figure 4.**
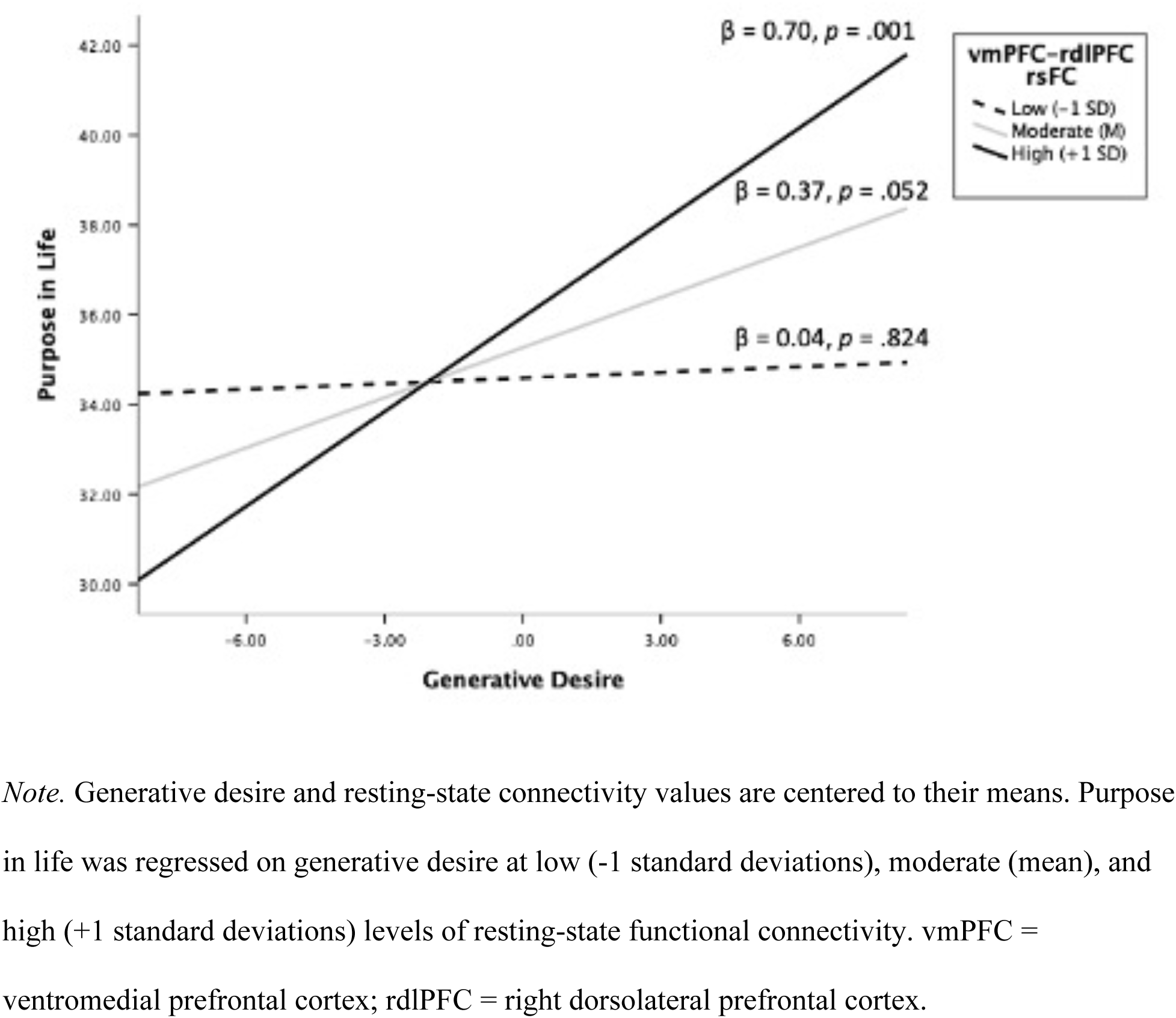
Interaction Between Generative Desire and Resting-State Functional Connectivity (rsFC) on Purpose in Life.

## 4. Discussion

The aim of the present study was to identify neurobehavioral factors influencing the relationship between generativity and purpose in life in older adults who are at risk for AD. This research builds upon prior work indicating an association between generativity and purpose in life in older adults (de St. Aubin, 2013). The current findings expand upon this by revealing that the availability of affectionate social support mediated the relationship between generative desire and purpose in life. Additionally, there was a significant association between generative desire and heightened vmPFC-rdlPFC and VS-precuneus rsFC. The strength of vmPFC-rdlPFC rsFC moderated the association between generative desire and purpose in life.

The behavioral mediation analysis revealed a positive association between generative desire and affectionate social support, which in turn was positively associated with purpose in life. The relationship between generative desire and affectionate social support aligns with the broaden-and-build theory, which posits that intrinsic motivation (i.e., generative desire) can augment positive emotions that allow individuals to build greater social resources, amplifying the availability of social support (Fredrickson, 2001). Thus, it is possible that older adults with higher generative desire cultivate more relationships that provide them with love and affection, resulting in greater life purpose. These results are also consistent with motivational lifespan theories, such as SST, that propose that as individuals age, they prioritize goals that optimize their socioemotional well-being. Given prior research showing that generativity is a modifiable psychological state (Gruenewald et al., 2016), the current results suggest that increasing generative desire could be a means for older adults to enrich their socioemotional state (i.e., their perceived love and affection from others), thereby enhancing older adults’ ability to experience a sense of direction and life purpose. Enhancing older adults’ purpose in life is vital given that it tends to decline in old age, partly due to life changes such as retirement and widowhood (Pinquart, 2002). Moreover, high purpose in life during aging has been shown to protect against cognitive decline and mental and physical health issues (Boyle et al., 2010; Pinquart, 2002).

The findings of vmPFC-rdlPFC and VS-precuneus rsFC being positively associated with generative desire can be understood in light of prior fMRI and neurostimulation studies. Our findings highlight how generative desire is associated with functional coupling between task-negative and task-positive brain networks. The vmPFC is a region within the reward and default mode networks that is active during internally-directed, passive cognitive tasks such as reward valuation and self-transcendent processing (Bartra et al., 2013; Kang et al., 2018). Conversely, the dlPFC is a hub in the executive control network, involved in functions like planning and working memory during externally-directed tasks (Petrides, 2005). Similarly, the precuneus is a region of the default mode network, implicated in higher-order cognitive functions such as self-referential processing, episodic memory retrieval, and mental imagery (Cavanna & Trimble, 2006), while the VS is a subcortical region activated during reward anticipation and reinforcement learning (Bartra et al., 2013).

The enhanced vmPFC-rdlPFC in those with higher generative desire aligns with the default-executive coupling hypothesis of aging proposed by Spreng and Turner (2019) and the finding of diminished anti-correlation between medial prefrontal cortex and dlPFC in aging (Keller et al., 2013). According to this neurocognitive model of aging, there is an age-related shift in goal-directed cognition characterized by a decline in cognitive control abilities and an accumulation of semantic knowledge. The model suggests that the increased reliance on semantic knowledge in aging is associated with concurrent heightened executive control network activity (e.g., lateral PFC) and reduced suppression of the default mode network (e.g., medial PFC). This coupling has also been shown to extend to regions within the reward network during goal simulation. In an fMRI study by Gerlach et al. (2014), participants mentally simulated the process of achieving a goal and, subsequently, the experience of achieving that goal. They showed that simulating the goal achievement process resulted in enhanced functional connectivity between default mode and executive control network regions, whereas simulating the experience of achieving the goal involved enhanced connectivity between default mode and reward network regions. Taken together, these results imply that generative desire in older adults involves a heightened reliance on semantic knowledge rather than cognitive control processes when cultivating future goals. The default mode network may support older adults’ ability to access internal mental representations of themselves and the world when envisioning goals that promote the well-being of future generations.

In addition to goal setting, vmPFC-rdlPFC connectivity has also been observed in prosocial decision making. In one study, participants who underwent transcranial magnetic stimulation (TMS) to the rdlPFC rejected selfish options in favour of costly options that benefited others (Baumgartner et al., 2011). The prioritization of others over the self was associated with enhanced connectivity between vmPFC to rdlPFC. The authors concluded that the rdlPFC might exert top-down executive control over the vmPFC, thus preventing the impulse to act selfishly in favour of a prosocial alternative. These results highlight how enhanced vmPFC to rdlPFC rsFC in those with higher generative desire might support older adults’ ability to suppress self-serving impulses or tendencies, thereby facilitating the pursuit of prosocial, generative goals. This aligns with Erikson’s (1950) definition of generativity as the “ability to transcend personal interests to provide care and concern for younger and older generations.”

We additionally found that connectivity between the vmPFC and rdlPFC moderated the association between generative desire and purpose in life, with high rsFC (+1 SD above the mean) demonstrating the strongest, positive relationship between generative desire and purpose in life. This highlights how individual differences in brain connectivity might influence older adults’ ability to derive life purpose from generative goals. In particular, it underscores how there is a need to consider individual differences in order to develop the right interventions to enhance generativity and purpose in life for the right individual at the right time. Moreover, it demonstrates the potential for future therapeutics (e.g., rTMS) that increase vmPFC-rdlPFC rsFC to bolster the relationship between generative desire and purpose in life, particularly in older adults at risk of AD.

Nonetheless, the present study does have certain limitations. The brain imaging data were acquired two years prior to the collection of the behavioral data. Given that generativity and rsFC are impacted by age-related processes, this presents a putative temporal confound. Furthermore, this study is limited by the over-representation of women in the current sample. Thus, these findings should be replicated in diverse samples in order to strengthen the generalizability of the findings. While all participants in this study are at risk of AD due to a first-degree family history, no participants in this study were diagnosed with mild cognitive impairment (MCI) or early dementia. A potential avenue for future research would be to examine the presence of AD biomarkers (e.g., PET amyloid-tau) in the vmPFC, rdlPFC, VS, and precuneus of individuals with MCI or early dementia and their impact on generative desire over the disease course, thereby offering insights into the relationship between generativity, AD biomarkers, and cognitive functioning.

In summary, this study demonstrated that generative desire was associated with greater rsFC between the vmPFC and rdlPFC as well as the VS and precuneus, regions implicated in goal-directed cognition and prosociality. Moreover, vmPFC-rdlPFC rsFC and affectionate social support were found to moderate and mediate the relationship between generative desire and purpose in life, respectively. The results of the current study have practical implications for promoting the health and well-being of at-risk older adults. Interventions designed to enhance generativity and affectionate social support may be effective to increase purpose in life in older adults at risk for cognitive impairment (Gruenewald et al., 2016). Future interventions might increase their efficacy by implementing familial intergenerational strategies that enhance feelings of love, appreciation, and affection in older adults. Prior research has demonstrated that older adults in early stages of AD experience preserved socioemotional functioning (Sturm et al., 2013; Zhang et al., 2015), suggesting that such interventions might also be effective for those with early functional impairment. Finally, a deeper understanding of the brain basis of generativity is crucial to understand the cognitive processes and brain mechanisms that support healthy aging. These findings may guide the development of personalized interventions and circuit-based strategies to promote resilient aging.

## 6. Conflict of Interest

The authors have no conflicts of interest to disclose.

## 7. Funding

This work is supported by a National Sciences and Engineering Research Council of Canada (NSERC) Discovery Grant (DGECR-2022-00299) and a NSERC Early Career Researcher Supplement (RGPIN-2022-04496), NIH P30 AG048785, a QBIN award and a Fonds de Recherche Santé Quebec (FRSQ) Salary Award to MRG. This research was also undertaken thanks to funding from the Canada First Research Excellence Fund, awarded to the Healthy Brains, Healthy Lives initiative at McGill University to CSW.

### 8. Acknowledgments

We are extremely grateful for the research participants who dedicated their time to participating in this study. We would also like to thank all members of the Pre-symptomatic Evaluation of Experimental or Novel Treatments for Alzheimer Disease (PREVENT-AD) Research Group. Finally, we would like to thank all lab members who participated in data collection for this study, including Maggie Nguyen, Sarah Elbaz, Andréanne Powers, Nicolas Lavoie, and Shania Fock Ka Bao. Data, analytic methods, and study materials will be made available to other researchers upon request. This study was not preregistered.

## Notes

### Competing Interest Statement

The authors have declared no competing interest.

### Summary of Updates

15 additional participants were added to the behavioral analyses and 7 additional participants were added to the neuroimaging analyses. 3 authors were added who provided critical feedback to the first version of the manuscript.

