## Supplementary information for "The influence of generativity on purpose in life is mediated by social support and moderated by prefrontal functional connectivity in at-risk older adults"

### 1. MRI Preprocessing

The preprocessing of both functional and structural data was performed using the fMRIprep pipeline. First, the structural images underwent skull stripping using a Nipype implementation of the antsBrainExtraction.sh tool (ANTs), an atlas-based brain extraction workflow. Next, brain tissue was segmented with FSL fast and normalized to Montreal Neurological Institute (MNI) space using ANT's antsRegistration. Preprocessing for the resting-state functional images included head motion correction, realignment, slice timing correction, susceptibility distortion correction, co-registration to reconstructed structural images, and spatial normalization to standard space. For each BOLD run, the BOLD time-series was averaged to generate a reference volume. The BOLD reference was co-registered to the T1W reference using bbregister, implementing a boundary-based registration with six degrees of freedom. Head motion parameters with respect to the BOLD reference (one rigid-body transformation, three rotations, and three translations) were estimated with FSL's mcflirt. Then, the rigid-body transformation was applied to re-sample the BOLD time series onto their original, native space. Transformations were concatenated to map the BOLD image to MNI standard space. Potential confounds were estimated, including the mean global signal, mean tissue signal class, tCompCor, aCompCor, Framewise Displacement, and DVARS. Volumes with framewise displacement above 0.9 mm and/or global blood-oxygen-level-dependent (BOLD) signal changes above 3 standard deviations were flagged as motion outliers. All dummy scans were removed prior to any additional analyses.

Denoising was performed on the fMRIprep outputs using Tedana. The signal across echoes were combined using a weighted average, and then normalized across echoes. A time series for the optimally combined echoes was then extracted. Principal component analysis was applied to the optimally combined data to separate the BOLD signal from thermal noise. Then, independent component analysis denoising was performed. This step uses a TE-dependence model to classify principal component analysis components as BOLD or non-BOLD while removing motion and physiological noise. Finally, the functional data were smoothed using a full-width half-maximum kernel of 6 mm in the CONN toolbox (Whitfield-Gabrieli & Nieto-Castanon, 2012).

**2. The TestMyBrain (TMB) Digital Neuropsychology Toolkit (Singh et al., 2021)**

The TMB Digital Neuropsychology Toolkit is a reliable and valid tool used for remotely administered neuropsychological evaluations. It includes a variety of computerized tests that assess working memory, attention, processing speed, memory, executive functioning, and reasoning. The tests used in the current study are described below:

*Simple Reaction Time.* This test measures basic psychomotor response speed. Participants are presented with red squares that say “STOP!” and green squares that say “GO!” and are instructed to press a key on their keyboard (or tap the screen) whenever a green square appears. Scores for each participant reflect their median reaction time.

*Choice Reaction Time.* This test measures the speed of response selection and attention. On each trial, participants are presented with 3 arrows, with one arrow having a different colour from the rest. Participants are asked to indicate the direction of the arrow that is a different colour from the rest by pressing the “x” key (to indicate left) or the “m” key (to indicate right). Scores for each participant reflect their median reaction time on correctly answered trials.

*Gradual Onset Continuous Performance Test.* This test measures sustained attention, response inhibition, and cognitive control. The participant is presented with images of cities that rapidly transition into other images or cities or images of mountains. Images of mountains appear 10-20% of the time. The participant is instructed to press a key when a city image appears and to not press anything when a mountain image appears. Discrimination scores reflect how accurately the participant was able to respond to mountains and cities, with higher scores indicating greater accuracy. The impulsivity score reflects how quickly and impulsively the participant responded, with higher scores indicating greater impulsivity.

*Matrix Reasoning.* This test is a measure of perceptual reasoning. On each trial, participants view a matrix of images with one image missing. They are asked to select the image (out of five response options) that best completes the pattern, based on a logical rule. Trials gradually increase in difficulty. Scores reflect the number of correct responses out of 36.

*Digit Span (Forward and Backward).* This test measures working memory and attention. Participants are presented with a series of digits for 1000 milliseconds each. In forward digit span, participants are asked to recall the sequence in the same order by typing the number series on the keyboard. In backward digit span, participants are asked to recall the sequence in the reverse order by typing the sequence on their keyboard. In both tests, the number series begins with two numbers and increases to a maximum of 11 numbers. Participants complete two trials at each span length and if at least one trial is answered successfully, the span length is increased by one number. If both trials with the same span length are answered incorrectly, the task is discontinued. Scores on both tests are the longest span length where at least one trial was answered successfully.

*Visual Paired Associates.* This test measures episodic memory. Participants are presented with 24 pairs of images and are told that they will later be tested on which images were paired together. After all image pairs are presented, there is a delay of approximately 2.5 minutes where participants complete the Choice Reaction Time test. Then, participants are presented with one image from each of the pairs that were previously presented and are asked to select which image was presented with it. Participants are presented with five response options. Scores reflect the number of images that were correctly identified, out of 24.

*Digit Symbol Matching.* This test is a measure of processing speed. On all trials, participants are presented with nine symbols that are paired with the digits 1, 2, or 3. On each trial, one symbol is presented above these pairings and participants are asked to press the key of the corresponding digit as quickly as possible. Scores reflect the total number of correct responses in 90 seconds.

#### 3. Seed-to-Voxel Connectivity Maps

**Figure 1.** Group-level whole brain correlations with the ventromedial prefrontal cortex. Results are corrected for multiple comparisons and overlaid on a standard template.

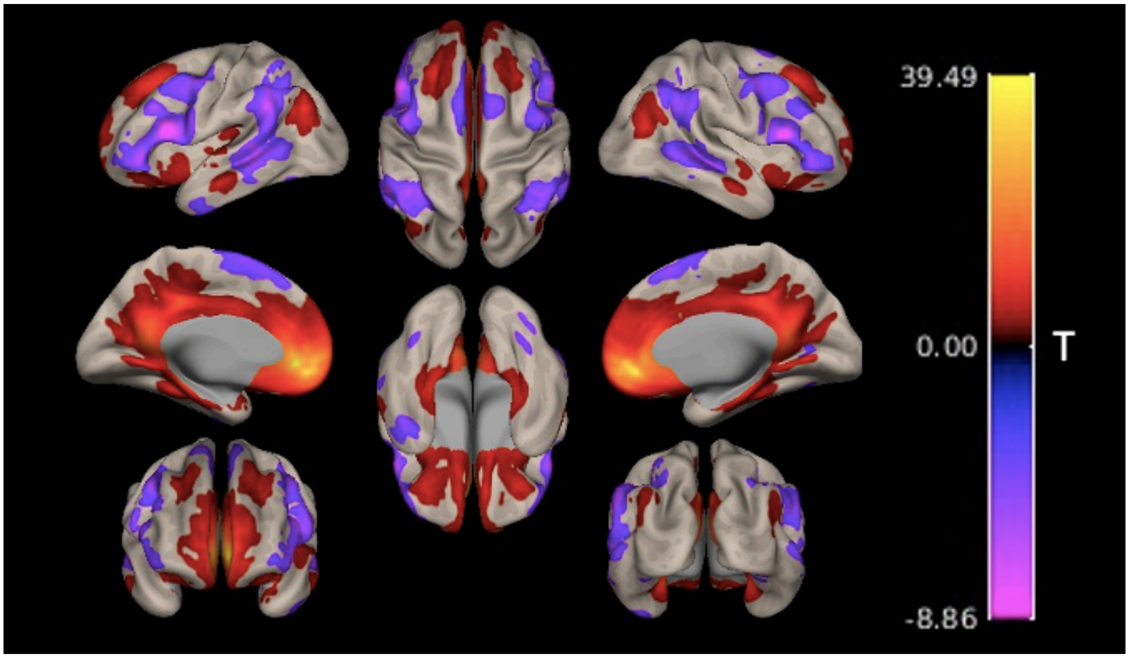


**Figure 2.** Group-level whole brain correlations with the ventral striatum. Results are corrected for multiple comparisons and overlaid on a standard template.

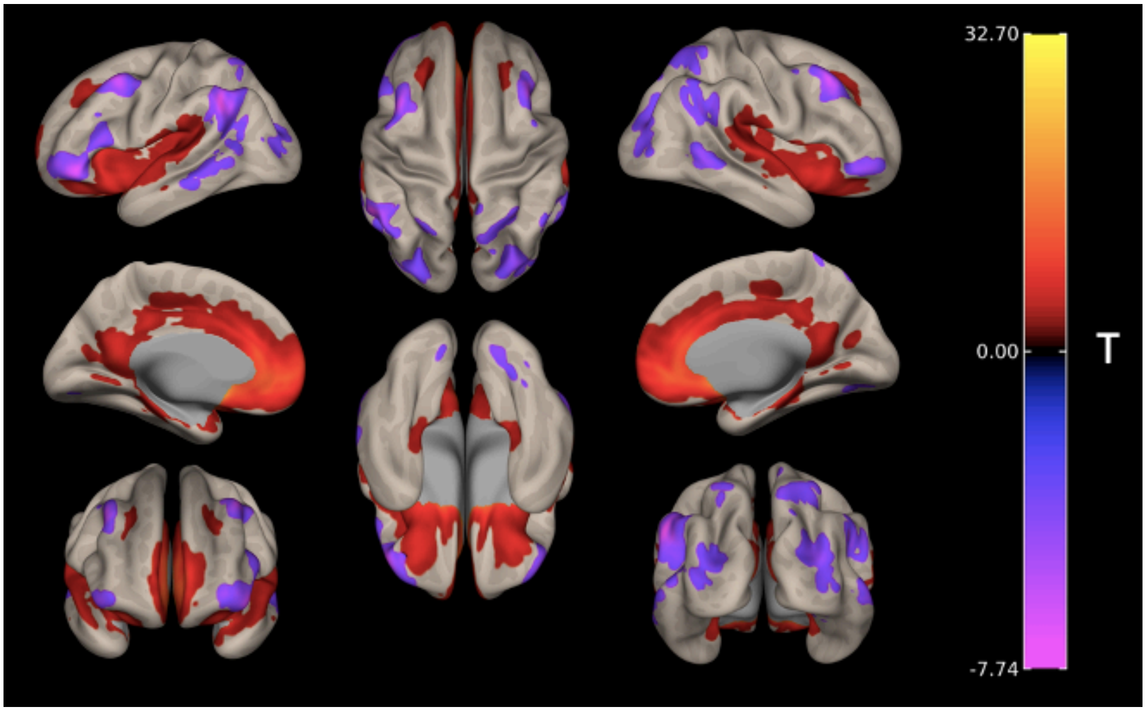


Seed-to-voxel analyses were conducted to identify brain regions demonstrating correlated and anticorrelated activity with the vmPFC and VS ROIs. Figure 1 demonstrates the expected vmPFC functional connectivity to regions of the reward (i.e., caudate, putamen, nucleus accumbens, hypothalamus, hippocampus, amygdala, orbitofrontal cortex), frontoparietal (e.g., precuneus, frontal poles), and default mode networks (i.e., cingulate gyrus, superior frontal gyrus, medial frontal cortex), as defined by Yeo et al.'s (2011) seven-network parcellation. This ROI was also anticorrelated with certain regions of the default mode network (e.g., inferior parietal lobule and superior and medial temporal cortex). Figure 2 shows the expected VS functional connectivity to the medial prefrontal cortex and anterior cingulate cortex as well as subcortical regions including the caudate, putamen, nucleus accumbens, hippocampus, and amygdala. This ROI was anticorrelated with regions of the lateral frontal cortex (frontal pole), occipital cortex (lateral occipital cortex, intracalcarine cortex, occipital pole) and parietal cortex (angular gyrus, supramarginal gyrus, superior parietal lobule).
